# Eye proteome of *Drosophila melanogaster*

**DOI:** 10.1101/2023.03.04.531088

**Authors:** Mukesh Kumar, Canan Has, Khanh Lam-Kamath, Sophie Ayciriex, Deepshe Dewett, Mhamed Bashir, Clara Poupault, Kai Schuhmann, Oskar Knittelfelder, Bharath Kumar Raghuraman, Robert Ahrends, Jens Rister, Andrej Shevchenko

## Abstract

The *Drosophila melanogaster* eye is a popular model to elucidate the molecular mechanisms that underlie the structure and function of the eye as well as the causes of retinopathies. For instance, the *Drosophila* eye has been used to investigate the impacts of ageing and environmental stresses such as light-induced damage or dietary deficiencies. Moreover, large-scale screens have isolated genes whose mutation causes morphological and functional ocular defects, which includes key components of the phototransduction cascade. However, the proteome of the *Drosophila* eye is poorly characterized. Here, we used GeLC-MS/MS to quantify 3516 proteins he adult *Drosophila melanogaster* eye and provide a generic and expandable resource for further genetic, pharmacological, and dietary studies.

## Introduction

The *Drosophila melanogaster* is an established model organism to elucidate the molecular mechanisms that underlie the structure and function of the eye and how their disruption causes retinopathies [1–6]. The adult compound eye is a highly repetitive structure that consists of about 800 units that are called ommatidia. Each ommatidium contains eight photoreceptor neurons (R1-R8) that express different light-sensing Rhodopsins and phototransduction proteins in specialized light-sensing compartments [7–9], which depends on vitamin A [10, 11]. The photoreceptors are surrounded by accessory cells that include cone, pigment, and mechanosensory bristle cells [12].

A genetic toolkit available in *Drosophila* allows the generation of loss-of-function clones specifically in the eye [13–15]. Large-scale screens isolated genes whose mutation causes morphological and functional ocular defects [16–19]. This approach led to the identification of signaling pathways that mediate major developmental processes and unraveled the components of the phototransduction cascade[20]. For instance, a crucial discovery was the identification of light-activated <u>T</u>ransient <u>R</u>eceptor <u>P</u>otential (TRP) channels [21] that founded a conserved superfamily of cation channels with diverse functions ranging from mediating the responses to various sensory stimuli [22] to immune responses [23]. Moreover, TRP channels have been implicated in various human diseases including cancer and neurodegenerative disorders [24, 25]. Lastly, the *Drosophila* eye has also been used as a model to investigate the impacts of ageing [26] and environmental stresses such as light-induced damage [27–29] or diets that lack essential nutrients [10, 30, 31].

Despite these important discoveries, *Drosophila* eye proteome is poorly characterized. While several resources for gene expression in the adult eye are available, such as transcriptomes of whole eyes, isolated photoreceptor nuclei, or single cells [26, 32–35], transcript levels appear to only weakly correlate with protein expression levels [36–38]. Yet, there are very few resources for the ocular proteome or the molar abundances of proteins that are expressed in the eye [10, 39, 40]. Here, we used GeLC-MS/MS to quantify 3516 proteins in the eye of adult *Drosophila* animals. Approximately 30% of them were identified as membrane-related proteins and 16% have at least one transmembrane domain. We also quantified the absolute (molar) abundances of a functionally related set of proteins that is critical for phototransduction and photoreceptor morphology. Taken together, we provide a quantitative and expandable resource for further genetic, pharmacological, and dietary studies.

## Results and Discussion

A schematic of the experimental workflow used in this study is shown in **Figure 1**. *Drosophila melanogaster* were raised from the embryonic to the adult stage on ‘standard’ lab food (SF) under a 12h light/12h dark cycle at 25°C (for details, see Materials and Methods). Three to four days-old male flies were collected and used for all the experiments described below.

**Figure 1:**
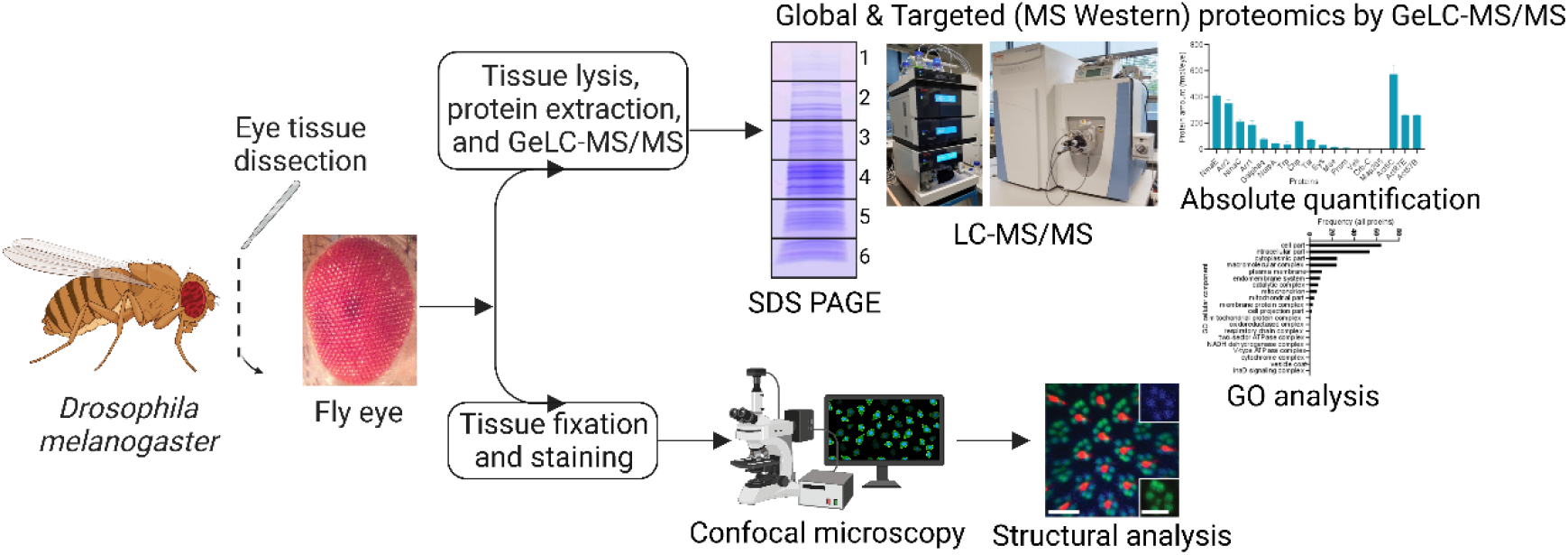
Schematic of the workflow used in this study. Experimental workflow for cataloging the proteome of the *Drosophila melanogaster* compound eye and absolute (molar) quantification of major proteins that are essential for photoreceptor structure and function. Schematics were created with BioRender. For details, see text.

The nutrient-rich SF led to normal morphology of the compound eye (**Figure 2A**) and wild type ommatidia with six rhabdomeres of the rod-equivalent outer rhabdomeres arranged in a trapezoid shape around the rhabdomeres of the cone-equivalent inner photoreceptors (**Figure 2B**). Moreover, we detected a wild-type expression pattern of the major Rhodopsin Rh1 in the outer rhabdomeres (**Figure 2B**), which suggests that SF supports wild type visual pigment formation and is vitamin A-sufficient [10, 41].

**Figure 2:**
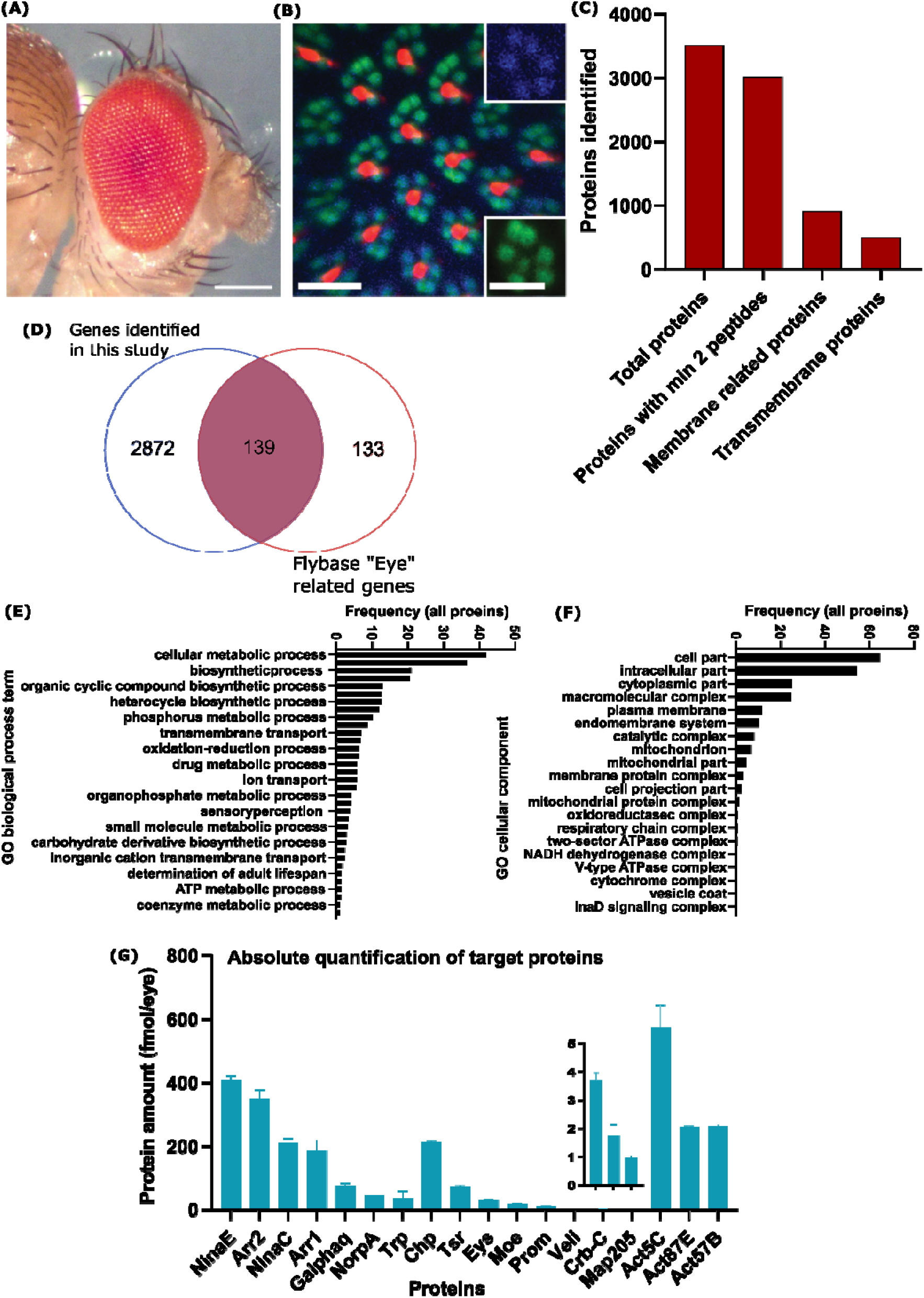
The eye structure and proteome of wild-type *Drosophila melanogaster* raised on ‘standard’ lab food. **(A)** Side view of the compound eye of a male wild-type fly that was raised on ‘standard’ lab food (SF). Scale bar, 10 μm. **(B)** Adult retina confocal cross-sections of male wild type fly raised on SF. Seven F actin-rich (Phalloidin, green) rhabdomeres are visible in each unit eye; Rh1 (blue inset) is expressed in the rhabdomeres of ‘outer’ photoreceptors and Rh6 (red) in the rhabdomeres of ‘inner’ photoreceptors. Scale bars, 10 μm (insets, 5 μm). **(C)** Graph showing the total number of identified proteins, the number of proteins that were identified with at least two peptides per protein, as well as membrane-related and plasma membrane proteins. (**D**) Protein-coding genes that are expressed in the eye identified in this study (blue) compared with ‘eye’-related genes annotated in the FlyBase database (red; from www.flybase.org). (**E**) Enrichment analysis for the GO term ‘cellular component’ of all quantified (iBAQ) proteins. (**F**) GO enrichment analysis assigned identified membrane proteins to 20 cellular components. (**G**) MS Western quantification of the absolute (molar) abundances of proteins that play a major role in photoreceptor morphology or phototransduction.

To characterize the adult eye proteome, we analyzed protein extracts by label-free GeLC-MS/MS quantitative proteomics by. We quantified a total of 3516 proteins (3017 of them with at least two peptides) (**Figure 2C**); the complete list of quantified proteins is provided in **Table 1**. According to FlyBase database (FB2022_02, released March 29, 2022), 2872 proteins were not attributed to eye tissues previously (**Figure 2D and Supplementary Table 2**). Approximately 30% of all quantified proteins (910/3017) are membrane-related proteins and 16% of them (502/3017) had at least one transmembrane domain (**Figure 2C**). To quantify and classify the identified proteins, we used intensity-based absolute quantification (iBAQ); based on the iBAQ value, we performed functional annotation and classification by Gene Ontology (GO) enrichment analysis. 34 terms that describe different biological processes were assigned to the proteins identified in this study, of which cellular metabolic, multicellular organismal, and biosynthetic process represented the major processes (**Figure 2E**). A GO enrichment analysis of these membrane proteins assigned them to 20 cellular components (**Figure 2F**).

Next, we used MS Western to quantify absolute (molar) abundances of proteins that play crucial roles in eye photoreceptor structure and function. MS Western method that allows the multiplexed, antibody-free and label-free molar quantification of user-selected proteins [42]. We found that the visual pigment Rhodopsin Rh1 (NinaE) was the most abundant phototransduction-related protein (~409 fmoles/eye) and the next three most abundant proteins are all involved in the termination of the light response (**Figure 2G**): the major visual arrestin Arr2 (~350 fmoles/eye), the myosin NinaC (210 fmoles/eye), and the other visual arrestin Arr1 (~185 fmoles/eye). The downstream phototransduction proteins activated by Rh1 [20]: G protein alpha subunit Galphaq (~76 fmoles/eye), NorpA (~45 fmoles/eye), and the major light-sensitive cation channel Trp (~36 fmoles/eye) were ca 10-fold less abundant than Rh1. (**Figure 2G**).

We also used MS Western to compare the molar abundances of proteins that are required for photoreceptor morphology and maintenance [43]. The molar abundances of the structural proteins were consistent with their spatial expression pattern (broad or restricted) in the eye: the cell adhesion molecule Chaoptin (Chp) (~212 fmoles/eye), which is required for the adhesion of the rhabdomeric microvilli and is broadly expressed throughout their perimeter [44, 45], was the most abundant protein (**Figure 2G**). Much less abundant were two structural proteins with more restricted expression patterns that are critical for rhabdomere separation [46], the secreted glycoprotein Eyes shut (~31 fmoles/eye) that is localized to the interrhabdomeral space (Eys, also called Spacemaker/Spam) [45] and the transmembrane protein Prominin (Prom) (~10 fmoles/eye) that is spatially restricted to the stalk membrane of the rhabdomeres as well as the tips of the microvilli [45]. The transmembrane protein Crumbs (Crb) (~2 fmoles/eye) that is essential for rhabdomere morphology was the least abundant morphology protein, which is consistent with its even more restricted expression exclusively in the stalk membrane [47].

Lastly, we quantified the molar abundances of the three Actins Act5C, Act87E, and Act57B that are expressed in the adult eye [40]. Act5C was expressed at even higher levels than Rh1 and also the most abundant actin (~573 fmoles/eye). The molar abundances of Act87E and Act57B were about half of those of Act5C, ~260 and ~262 fmoles/eye, respectively (**Figure 2G**). Lastly, the scaffolding protein and Crumbs complex member Veli/Lin-7 (~4 fmoles/eye) [48] and the actin depolymerization factor Twinstar (a homolog of Cofilin/ADF) (~73 fmoles/eye), showed significantly lower absolute abundances (**Figure 2G**).

## Conclusion

Taken together, our study provides a comprehensive catalogue of the proteome of adult *Drosophila melanogaster* eye. We envision that these data will be a useful resource for the scientific community that uses the *Drosophila* eye as a model to study visual signaling or the genetic and environmental stresses that cause various retinopathies.

## Material and Methods

### *Drosophila* stock keeping

The *Drosophila melanogaster* wild-type strain Oregon R was reared under a 12h light/12h dark cycle at 25°C. The flies were raised on ‘standard’ lab food (SF), which contained per liter: 8g agar, 18g brewer’s yeast, 10g soybean, 22g molasses, 80g cornmeal, 80g malt, 6.3ml propionic acid, and 1.5g Nipagin.

### *Drosophila* compound eye images

Adult male *Drosophila melanogaster* were immobilized with CO_2_ and embedded in a liquid agarose gel as previously described [10]. After solidification of the gel, the petri dish was placed under the Stemi 508 Trinoc microscope (Zeiss model #4350649030), and the fly head was adjusted with forceps such that one compound eye faced the microscope lens. Imaging was performed with an Axiocam 208 HD/4k color camera (model #4265709000) that was set to auto exposure and auto white balance. Pictures were processed with Fiji, Adobe Photoshop 2020, and Adobe Illustrator 2020 software.

### Confocal microscopy and immunohistochemistry of *Drosophila* photoreceptors

We dissected retinas of four days old flies as previously described [49]. Briefly, we fixed the retinas in 3.8% formaldehyde solution before removing the laminas and head cuticle. Retinas were incubated overnight in the primary antibody (mouse anti-Rh1 4C5, 1:10, from Developmental Studies Hybridoma Bank, University of Iowa) diluted in PBST (PBS + 0.3% Triton-X, Sigma). The next morning, retinas were washed three times with PBST. Retinas were then incubated overnight in secondary antibody diluted in PBST (1:800, Alexa Fluor 647-conjugated raised in donkey, Invitrogen) and Alexa Fluor 488-conjugated Phalloidin (1:100, Invitrogen). The next morning, three PBST washes followed. Retinas were mounted on a bridge slide using SlowFade (Molecular Probes) and imaged with a Zeiss LSM 8 confocal microscope. Raw images were processed with Fiji (https://imagej.net/software/fiji/) and then cropped and contrasted with Adobe Photoshop and Adobe Illustrator.

### Protein extraction and GeLC-MS/MS analysis

The compound eyes (n=40) were dissected (including the lamina neuropil) from the three to four days old flies raised on SF and placed in lysis buffer containing 150 nM NaCl, 1 mM EDTA, 50 mM Tris-HCl (pH7.5), 1 tablet Roche protease inhibitors, 0.2% w/v CHAPS, 0.1% w/v OGP, 0.7% v/v triton X-100, 0.25 μg/mL DNase and RNase. The samples were immediately snap frozen using liquid nitrogen and stored at −80°C or further processed. The eye tissues were homogenized and to the supernatant an equal volume of 2X SDS Laemmli sample buffer (SERVA Electrophoresis GmbH, Heidelberg, Germany) was added. The samples were heated at 80°C for 10-15 minutes and loaded on 4-20% 1D SDS PAGE. The protein bands were visualized by Coomassie Brilliant Blue staining. Each gel lane was cut into six gel slices, and each gel slice was co-digested with heavy isotope labeled CP02 and gel band containing 1pmol of BSA standard.

In-gel digestion was carried out as per the protocol described [50]. Briefly, the electrophoresed gel, rinsed with water, was stained with Coomassie Brilliant Blue R-250 for 10 minutes at RT and then destained with destaining solution (Water: Methanol: Acetic acid, 50:40:10 (v/v/v). The gel slice was excised as per the expected molecular weight of the proteins of interest and further cut into small pieces (~1 mm size). The gel pieces were then transferred into 1.5 ml LoBind Eppendorf tubes and further processed. The gel pieces were completely destained by ACN/Water, reduction was done by incubating the gel pieces with 10 mM Dithiothreitol at 56°C for 45 minutes. Alkylation was carried out with 55mM Iodoacetamide for 30 minutes in dark at RT. The reduced and alkylated gel pieces were washed with water/ACN and finally shrunk with ACN, ice-cold trypsin (10ng/μl) was added to cover the shrunk gel pieces and after 1hr of incubation on ice the excess trypsin (if any) was discarded. The gel pieces were then covered with 10mM NH4HCO3 and incubated for 12-15hrs at 37°C. The tryptic peptides were extracted using water/ACN/FA, dried using a vacuum centrifuge and stored at −20°C until use. The tryptic peptides were recovered in 5% aqueous FA and 5 μL were injected using an autosampler into a Dionex Ultimate 3000 nano-HPLC system, equipped with a 300 μm i.d. × 5 mm trap column and a 75 μm × 15 cm Acclaim PepMap100 C18 separation column. 0.1% FA in water and ACN were used as solvent A and B, respectively. The samples were loaded on the trap column for 5 min with solvent A at a flow of 20 μL/min. The trap column was then switched online to the separation column, and the flow rate was set to 200nL/min. The peptides were fractionated using 180 min elution program: a linear gradient of 0% to 30% B delivered in 145 min and then B% was increased to 100% within 10 min and maintained for another 5 min, dropped to 0% in 10 min and maintained for another 10 min. Mass spectra were acquired using either LTQ Orbitrap Velos or Q Exactive HF mass spectrometer both from Thermo Fisher Scientific (Bremen, Germany). The Data Dependent Acquisitions (DDA) settings used for both the mass spectrometers are provided in **Supplementary Table 1.**

### Data processing for protein identification and quantification

Mascot v2.2.04 (Matrix Science, London, UK) was used for peptide identifications against the custom-made database containing the sequence of the target protein, to which sequences of human keratins and porcine trypsin were added. For eye proteome analysis, the *Drosophila* reference proteome database from UniProt was used. The database searches were performed with the following mascot settings: precursor mass tolerance of 5 ppm; fragment mass tolerance of 0.6 Da and 0.03Da for the Velos and QE-HF data respectively; fixed modification: carbamidomethyl (C); variable modifications: acetyl (protein N-terminus), oxidation (M); Label: 13C (6) (K), Label: 13C (6) 15N (4) (R), 2 missed cleavages were allowed. Progenesis LC-MS v4.1 (Nonlinear dynamics, UK) was used for the peptide feature extraction and the raw abundance of identified peptide was used for absolute quantification. MaxQuant v1.5.5.1 and Perseus v1.5.5.3 was used for label-free quantification and subsequent statistical analysis. MaxQuant analysis was done with default settings.

## Acknowledgements/funding

This publication was supported by the National Eye Institute of the National Institutes of Health under Award Number R01EY029659 to J.R. The content is solely the responsibility of the authors and does not necessarily represent the official views of the National Institutes of Health. The funders had no role in study design, data collection and analysis, decision to publish, or preparation of the manuscript. Work in AS laboratory was funded by MPI of molecular cell biology and genetics.

